# Molecular architecture of the DNA‐binding sites of the P‐loop ATPases MipZ and ParA from *Caulobacter crescentus*

**DOI:** 10.1101/766287

**Authors:** Yacine Refes, Binbin He, Laura Corrales-Guerrero, Wieland Steinchen, Gaël Panis, Patrick H. Viollier, Gert Bange, Martin Thanbichler

## Abstract

Two related P-loop ATPases, ParA and MipZ, mediate the spatiotemporal regulation of chromosome segregation and cell division in *Caulobacter crescentus*. Both of these proteins share the ability to form dynamic concentration gradients that control the positioning of regulatory targets within the cell. Their proper localization relies on their nucleotide-dependent cycling between a monomeric and a dimeric state, driven by interaction with the chromosome partitioning protein ParB, and on the ability of the dimeric species to associate non-specifically with the nucleoid. In this study, we use a combination of genetic screening, biochemical analysis and hydrogen/deuterium exchange mass spectrometry to identify the residues mediating the interaction of MipZ with DNA. Our results show that the DNA-binding activity of MipZ relies on a series of positively charged and hydrophobic residues lining both sides of the dimer interface. MipZ thus appears to associate with DNA in a sequence-independent manner through electrostatic interactions with the DNA phosphate backbone. In support of this hypothesis, chromatin immunoprecipitation analyses did not reveal any specific target sites *in vivo*. When extending our analysis to ParA, we found that the architectures of the MipZ and ParA DNA-binding sites are markedly different, although their relative positions on the dimer surface and their mode of DNA binding are conserved. Importantly, bioinformatic analysis suggests that the same principles apply to other members of the P-loop ATPase family. ParA-like ATPases thus share common mechanistic features, although their modes of action have diverged considerably during the course of evolution.

**SIGNIFICANCE:** ParA-like P-loop ATPases are involved in a variety of cellular processes in bacteria, including chromosome and plasmid segregation, chemoreceptor and carboxysome positioning, and division site placement. Many members of this large protein family depend on the ability to bind non-specific DNA for proper function. Although previous studies have yielded insights in the DNA-binding properties of some ParA-like ATPases, a comprehensive view of the underlying mechanisms is still lacking. Here, we combine state-of-the-art cell biological, biochemical and biophysical approaches to localize the DNA-binding regions of the ParA-like ATPases MipZ and ParA from *Caulobacter crescentus*. We show that the two proteins use the same interface and mode of action to associate with DNA, suggesting that the mechanistic basis of DNA binding may be conserved in the ParA-like ATPase family.

## INTRODUCTION

ParA-like ATPases play a central role in the subcellular organization of bacterial cells. They are evolutionarily related to the P-loop GTPase superfamily (1, 2) and characterized by a conserved nucleotide-binding pocket containing a deviant Walker A motif (known as the P-loop) and a Walker B motif, required for ATP binding and hydrolysis (3). The members of this family share significant sequence and structural similarity and dimerize in an ATP-dependent manner (4, 5). Moreover, they typically show a dynamic behavior *in vivo* that often involves changes in their localization patterns during the course of the cell cycle (6–10). Many ParA-like ATPases act as spatial regulators that orchestrate cellular processes by controlling the subcellular positioning of macromolecular structures (7, 11). The prototypical ParA homologs, for instance, mediate the partitioning of sister chromosomes and plasmids during cell division (9, 12). Several other members of this family (MipZ, MinD, PldP and PomZ) ensure proper division site placement by controlling the polymerization of the cell division protein FtsZ into the cytokinetic FtsZ ring (6, 8, 13–17). Other functions of ParA-like ATPases include carboxysome segregation (McdA) (18), the positioning of chemoreceptor clusters (ParC and PpfA) (19, 20) and DNA translocation during conjugation (21).

*Caulobacter crescentus* possesses two well-characterized P-loop ATPases, ParA and MipZ, which work together to coordinate chromosome segregation and cell division in this species. The functions of these two proteins are closely linked through their common interaction partner ParB, a DNA-binding protein that recognizes a cluster of centromere-like sites (*parS*) in the origin-proximal region of the chromosome (13, 22–24). At the start of the cell cycle, the origin region is anchored at the old pole through interactions between ParB and the polarity factor PopZ, with traces of FtsZ from the previous division event at the new pole (25, 26). At the beginning of S-phase, the *parS* cluster is duplicated, and one of the ParB•*parS* complexes is rapidly moved to the opposite pole and tethered to a new patch of PopZ, in an active process driven by ParA (6, 27–29). MipZ interacts dynamically with the two pole-associated ParB complexes and thus forms a bipolar gradient, with its concentration being highest at the poles and lowest at the cell center (6, 13). Since it acts as an inhibitor of FtsZ polymerization, the arrival of the moving ParB•*parS*•MipZ complex leads to the disintegration of the polar FtsZ assembly, and the released FtsZ molecules, along with newly synthesized ones, assemble into a Z-ring at midcell (6).

Both ParA and MipZ establish dynamic polar gradients, based on their ability to alternate between an ADP-bound monomeric and an ATP-bound dimeric state with distinct interaction patterns and diffusion rates, in a manner controlled locally by the ParB•*parS* complexes. Although the ATPase cycles of the two proteins are based on similar principles, differences in the regulatory effect of ParB lead to clearly distinct localization behaviors. In the case of ParA, ParB acts as an ATPase-activating protein. ParA-ATP dimers remain associated with the nucleoid until they interact with a moving ParB•*parS* complex, which triggers their disassembly into monomers that are released from the DNA (19, 27, 29–33). Repeated cycles of ParB binding and ATP hydrolysis thus lead to a gradual shortening of the ParA gradient, thereby promoting the directional movement of ParB across the nucleoid towards one of the cell poles. For MipZ, by contrast, the polar ParB•*parS* complexes serve as catalysts that recruit monomers and facilitate their dimerization, possibly by increasing their local concentration at the cell poles (6). Newly formed MipZ dimers have DNA-binding activity and are retained in the polar regions through association with the nucleoid (6, 13). Their lifetime is limited only by the low intrinsic ATPase activity of MipZ, which eventually generates monomers that return to the ParB•*parS* complexes, thereby restarting the cycle. As a result, MipZ forms stable bipolar gradients whose minimum at the cell center marks the site of cell division.

Various P-loop ATPases were shown to interact with the nucleoid to slow down their diffusion and thus enable the maintenance of subcellular protein gradients (13, 18, 19, 34). Their DNA-binding activity was suggested to be mediated by positively charged amino acids that are exposed on their surface (4, 20, 35, 36). Consistent with this idea, *in vivo* and *in vitro* studies of the *Bacillus subtilis* ParA homolog Soj (*Bs*ParA) identified two surface-exposed arginine residues that are essential for its interaction with DNA (35). Residues homologous to these arginines were later also implicated in the DNA-binding activity of PpfA from *Rhodobacter sphaeroides* and PomZ from *Myxococcus xanthus* (17, 20). More evidence for the importance of positively charged amino acids has recently come from the crystal structures of ParA-DNA complexes formed by ParA homologs from *Helicobacter pylori* (*Hp*ParA) (37) and *Sulfolobus solfataricus* pNOB8 (38), which each identified several surface-exposed lysine residues that are in direct contact with the DNA ligand. However, the binding interface pNOB8 ParA was different from that of its *H. pylori* homolog or *R. capsulatus* PpfA (38), suggesting that ParA proteins could have evolved different modes of DNA binding.

So far, the precise location of the DNA-binding interface of MipZ has remained unknown, because the residues contacting DNA in other ParA-like ATPases are not conserved in this protein. In this study, we identify and characterize nine surface-exposed residues surrounding the dimer interface of MipZ that are critical for DNA binding. Most of them are positively charged, suggesting that MipZ interacts with DNA non-specifically by contacting the phosphate backbone. Consistent with this notion, ChIP-Seq analysis reveals that MipZ interacts with a large number of chromosomal sites without a clear preference for a specific consensus sequence. Mapping the DNA-binding residues of *C. crescentus* ParA (*Cc*ParA), we further demonstrate that MipZ and ParA bind DNA in similar, positively charged regions, although the nature of the interacting residues varies considerably. Thus, while the biological functions of ParA-like ATPases have diverged significantly during the course of evolution, the general principle underlying their DNA-binding activity may be conserved.

## RESULTS

### Identification of DNA-binding residues on the MipZ surface

Our knowledge of the determinants mediating the DNA-binding activity of P-loop ATPases is largely based on studies of canonical ParA homologs. For this group of proteins, two distinct modes of DNA binding, mediated by different regions of the dimer surface, have been reported (37, 38). Moreover, the residues identified in previous studies are not universally conserved among DNA-binding members of the ParA-like ATPase family. To clarify which regions of MipZ mediate the interaction with DNA, we devised a reverse genetics approach. Taking advantage of the crystal structure of the MipZ dimer (PDB ID: 2XJ9) (13), we selected a variety of charged or bulky hydrophobic residues that are exposed on the MipZ surface (**Fig. 1A**) and exchanged them for alanine. The resulting 51 single-mutant MipZ variants were then analyzed for function *in vivo*. To this end, the mutant alleles were fused to *eyfp* (encoding yellow fluorescent protein) and expressed from an inducible promoter in cells depleted of the native MipZ. Subsequently, the morphology of the strains and the localization patterns of the mutant proteins were determined by microscopy. As a reference, this analysis also included a wild-type MipZ-eYFP fusion, a monomeric variant (K13A) unable to interact with DNA and FtsZ, and a constitutively dimeric variant (D42A) that is locked in the DNA-binding and FtsZ-inhibitory state (6, 13).

**Figure 1.**
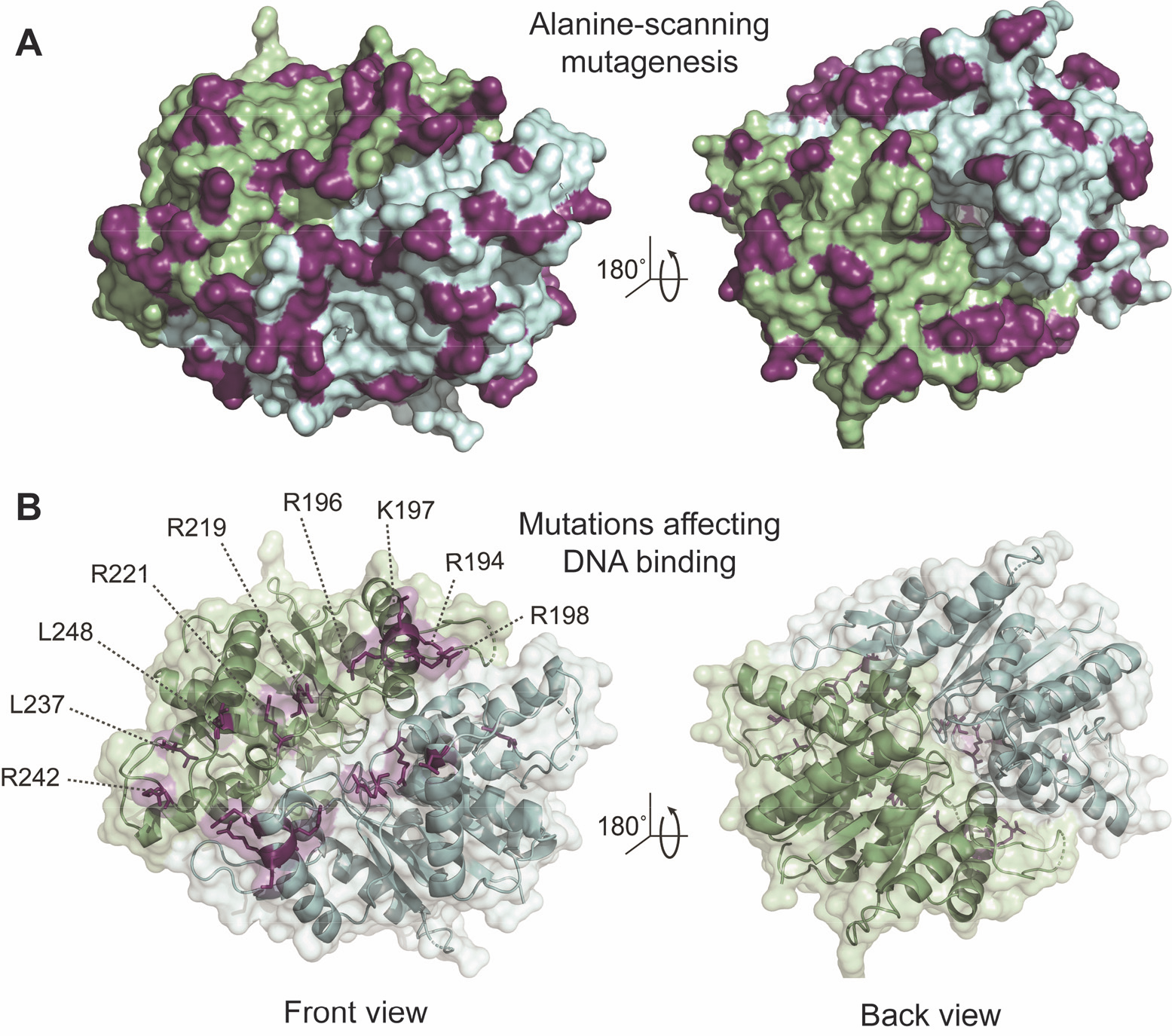
Identification of residues critical for DNA binding by MipZ. (**A**) Surface representation of the MipZ dimer structure (PDB ID: 2XJ9) with the 51 amino acids selected for alanine-scanning mutagenesis marked in purple. The two subunits are depicted in green and blue, respectively. (**B**) Cartoon representation of the MipZ dimer structure with nine candidate amino acids possibly involved in DNA binding shown in purple. A surface representation of the structure is shown in the background. For clarity, the ATP molecules and Mg^2+^ ions sandwiched between the two subunits are not shown.

As shown previously (6, 13), cells producing wild-type MipZ-eYFP displayed normal morphologies and the characteristic unipolar or bipolar gradient, indicating that the fusion protein was fully functional (**Fig. S1 and S2**). Expression of the K13A variant yielded a mixture of short and elongated cells as well as a high level of uniform cytoplasmic fluorescence, combined with distinct foci reflecting the positions of the ParB•*parS* complexes. The dimeric variant (D42A), by contrast, gave rise to filamentous cells showing patchy, nucleoid-associated fluorescence throughout the cell (**Fig. S1 and S2**).

DNA binding restrains the diffusion of MipZ and is a prerequisite for gradient formation (6). MipZ variants that fail to interact with DNA should still be able to associate with ParB. However, the dimers released from ParB should no longer be retained in the vicinity of the ParB•*parS* complexes but freely diffuse within the cytoplasm. As a consequence, they should inhibit cell division throughout the cell, leading to cell filamentation. Based on this hypothesis, we searched for mutant strains that still formed ParB-associated foci but displayed elevated levels of cytoplasmic fluorescence, combined with an increase in cell length. A total of nine strains showed the predicted phenotype (**Fig. 1B, S1 and S2A**). The stability of the different MipZ-eYFP variants was verified by immunoblot analysis (**Fig. S3A**). Expression of the R194A, R196A, R198A or R221A variants led to severe filamentation, whereas strains producing the K197A, R219A, L237A, R242A or L248A variants showed a milder phenotype with a broad distribution of cell lengths (**Fig. S2A**). Notably, the R196A, R219A and L248A variants produced very faint ParB-associated foci, suggesting that they have an additional defect in ParB binding (**Fig. S2A**). An analysis of the subcellular distribution of the different MipZ-YFP variants confirmed a partial to complete loss of the gradient pattern (**Figure S2B**). Interestingly, the nine amino acids identified in the screen are mostly positively charged and grouped in the same surface region to form an extended patch lining the dimer interface (**Figure 1B**).

### Diffusional properties of MipZ variants with DNA-binding defects

Previous work has shown that MipZ dimers have a very low diffusion rate, likely due to their interaction with the nucleoid. Monomers, by contrast, are highly mobile because they are rapidly exchanged between ParB•*parS* complexes and able to move freely within the cytoplasm (13). To corroborate that the newly identified variants have a reduced DNA-binding activity *in vivo*, we investigated the diffusional properties of two representative proteins (K197A and R198A) in fluorescence recovery after photobleaching (FRAP) assays. When one of the polar foci was bleached in cells showing a bipolar gradient of wild-type MipZ-eYFP, fluorescence was recovered with a half-time (t_½_) of 9.4 ± 0.4 sec (**Fig. S4A**). The monomeric K13A variant, by contrast, showed considerably faster kinetics (t_½_ = 2.0 ± 0.2 sec), indicating largely unrestrained diffusion (**Fig. S4B**). Notably, intermediate recovery rates were measured for both the K197A (t_½_ = 5.6 ± 0.2 sec) and the R198A variants (t_½_ = 5.3 ± 0.8 sec) (**Fig. S4C-D**). These results suggest that the two proteins are indeed less stably associated with the nucleoid and thus able to diffuse more rapidly within the cell than the wild-type protein. However, they still appear to interact with DNA to some extent, suggesting that they still display some residual DNA-binding activity (as confirmed below).

### Biochemical characterization of the DNA-binding-deficient MipZ variants

To confirm that the phenotypes observed are indeed due to a defect in DNA binding, we overproduced and purified the nine MipZ variants selected in the mutant screen and analyzed their biochemical properties *in vitro* (**Fig. S3B**). Initially, we performed ATPase activity assays to determine whether the mutant proteins were still able to dimerize and hydrolyze ATP. In all cases, the ATP turnover rates were similar to those of the wild-type protein (**Fig. 2A**), confirming that their ATPase cycles were not affected by the mutations. By contrast, and as expected (6), the monomeric (K13A) and dimeric (D42A) variants showed very low activity levels.

**Figure 2.**
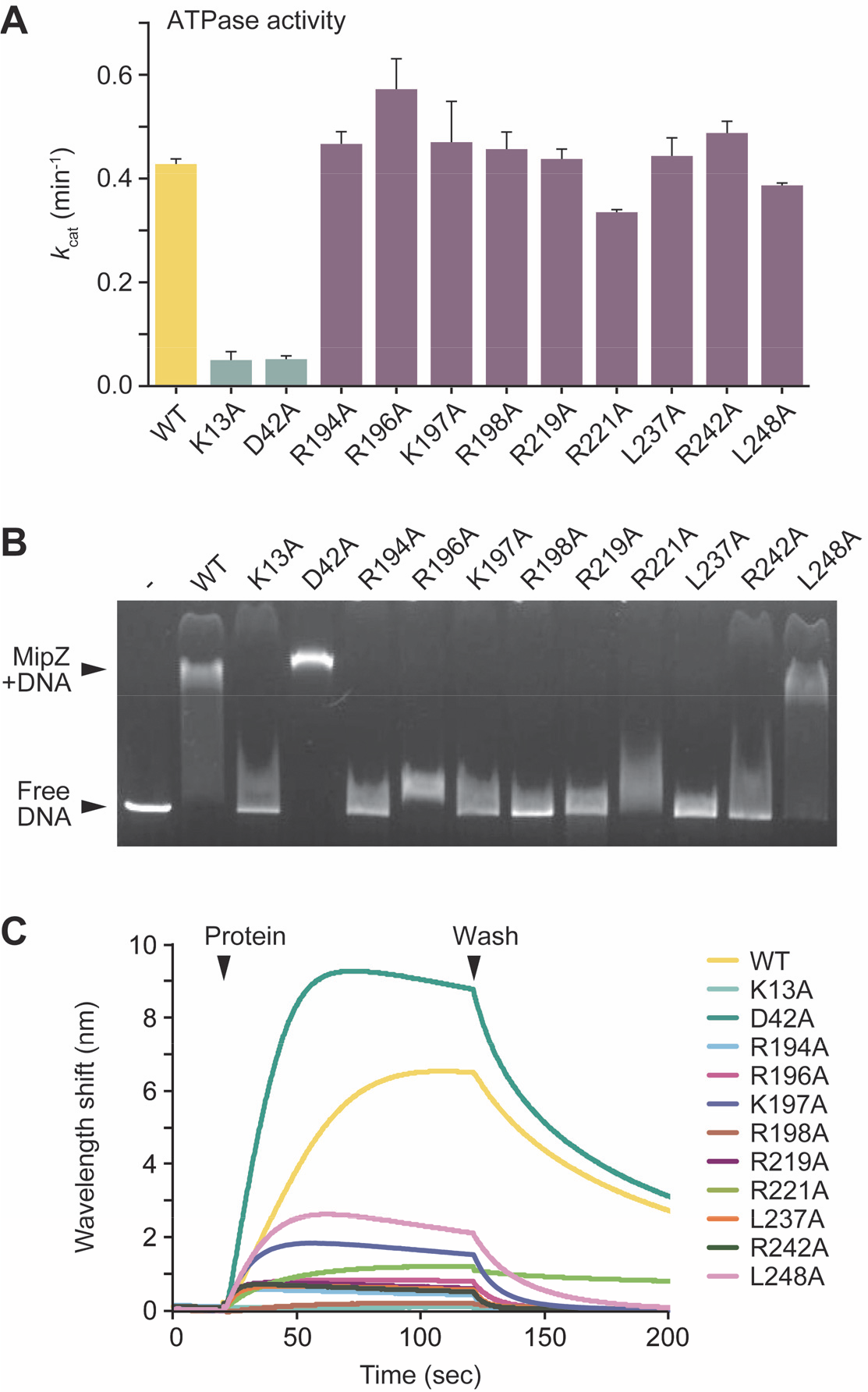
*In vitro* characterization of MipZ variants with DNA-binding defects. (**A**) ATPase activities of purified wild-type MipZ and the indicated mutant derivatives. MipZ (6 μM) was incubated with ATP (1 mM), and the rate of hydrolysis was determined with a coupled-enzyme assay. Shown are the average turnover numbers (*k*_cat_) (± SD) from at least three independent experiments. (**B**) Electromobility-shift assay analyzing the interaction of different MipZ variants with a linearized plasmid (pMCS-2). MipZ (WT) or the indicated variants (10 μM) were incubated with DNA (10 nM) and ATPγS (0.46 mM). Protein-DNA complexes were then separated from free DNA by standard agarose gel electrophoresis. (**D**) Biolayer interferometric analysis of the DNA-binding activity of different MipZ variants. A biotinylated dsDNA oligonucleotide (37.5 μM) was immobilized on the sensor surface and probed with MipZ (WT) or the indicated variants (4 μM) in the presence of ATPγS (1 mM). All interaction analyses were performed at least three times, and representative results are shown.

Next, we tested the DNA-binding capacity of the different variants using an electromobility shift assay (**Figure 2B**). For this purpose, the proteins were incubated with a linearized plasmid in the presence of the non-hydrolysable ATP analogue adenosine-5’-O-(3-thio)triphosphate (ATPγS), which locks MipZ in its dimeric state. Whereas the wild-type protein and the D42A variant strongly reduced the mobility of the DNA fragment during electrophoresis, most of the other mutant proteins showed hardly any binding activity. A slight shift was still observed for the R196A and R221A variants, while the L248A exchange hardly affected the behavior of MipZ in this assay. To corroborate these results, we additionally performed biolayer interferometric analyses. To this end, a double stranded (ds) 26 bp oligonucleotide was immobilized on a sensor chip and probed with wild-type MipZ or its mutant variants (**Fig. 2C**). The wild-type and D42A proteins again interacted strongly with DNA. The other mutant variants, by contrast, showed a moderate to severe decrease in the DNA-binding activity. An estimate of the changes in affinities was obtained by microscale thermophoresis experiments, in which a constant concentration of fluorescently (Cy3-) labeled dsDNA oligonucleotide was titrated with increasing concentrations of protein (**Fig. S5**). The data obtained enabled us to categorize the different MipZ variants into three groups: high-affinity binding proteins (*K*_D_ < 10 μM) (WT, D42A, L248A), low-affinity binding proteins (*K*_D_ ≫ 10 μM) (K13A, R196A, K197A, R219A, L237A, R242A) and proteins hardly showing any binding (R194A, R198A, R221A). Collectively, our *in vitro* results demonstrate that all MipZ variants selected in the mutant screen are defective in their interaction with DNA. Notably, the severity of the phenotypes observed for the different strains correlates with the DNA-binding affinities of the proteins they produce, underscoring the importance of DNA binding for proper MipZ function.

### MipZ binds DNA in a sequence-independent manner *in vivo*

Among the nine DNA-binding residues identified, seven are positively charged. The interaction of MipZ with DNA may therefore largely rely on electrostatic forces between positively charged MipZ residues and the negatively charged DNA phosphate backbone, as suggested for other ParA-like proteins (4, 20). Consistent with this hypothesis, MipZ indeed interacts non-specifically with DNA *in vitro* (13) (see also **Fig. 2**). However, it remained unclear whether it could have a preference for certain sequence motifs *in vivo*. To address this issue, we performed chromatin immunoprecipitation followed by high-throughput sequencing (ChIP-Seq) analysis on *C. crescentus* strains producing a wild-type MipZ-eYFP fusion or the corresponding K13A and D42A variants in place of the native MipZ protein (**Fig. 3**). As expected, DNA binding was undetectable for the monomeric K13A variant. The wild-type fusion also did not show any clear binding signals, likely because only a fraction of the protein is in the dimeric state and its ATPase activity greatly reduces the lifetime of DNA complexes. By contrast, a clear interaction with the nucleoid was observed for the D42A variant. Analysis of the precipitated DNA fragments revealed that it was associated with a large number of different loci. However, we did not observe distinct, localized signals, but broad peaks covering several kilobases of chromosomal DNA, suggesting that MipZ does not recognize defined sequence motifs. Interestingly, the extent of DNA binding was positively correlated with the GC content of the interacting chromosomal regions. However, *in vitro* binding studies did not provide any evidence for an intrinsic preference of MipZ for GC-rich DNA (**Fig. S6**). It has been shown that many nucleoid-associated proteins from *C. crescentus*, such as HU or GapR, are enriched in AT-rich chromosomal regions (39–41). Furthermore, AT-rich sequences are typically found at transcriptional promoters, which are heavily occupied by RNA polymerase or regulatory proteins. MipZ may therefore bind to DNA in a sequence-independent manner but, due to its moderate binding affinity, be restricted to less densely occupied, GC-rich regions.

**Figure 3.**
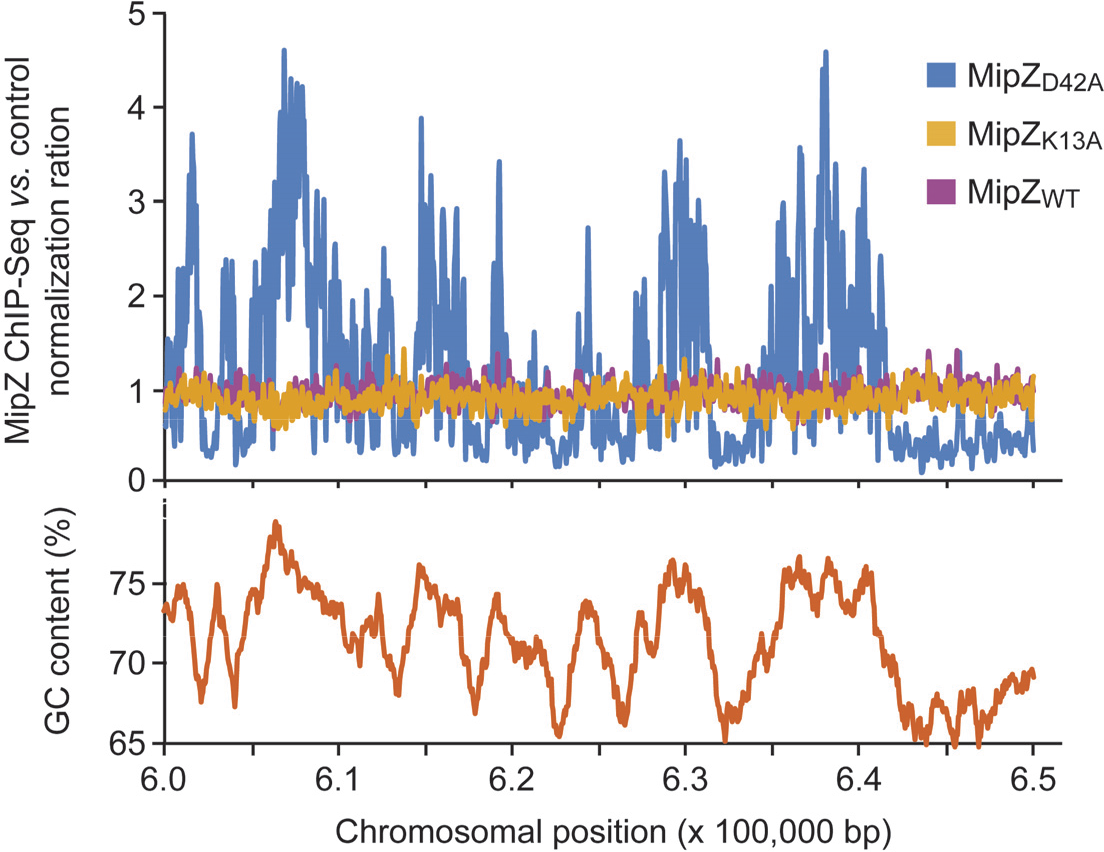
ChIP-seq analysis of the global distribution of chromosomal MipZ binding sites. *C. crescentus* strains producing a wild-type (BH64), monomeric (K13A; BH100) or dimeric (D42A; BH99) MipZ-eYFP fusion in place of the native protein were fixed with formaldehyde and subjected to ChIP-seq analysis with an anti-GFP antibody. A random 50 kb window of the chromosome is shown for visualization. Data were normalized using strain NA1000 as a background control.

### Direct detection of the MipZ DNA-binding interface by HDX analysis

Using reverse genetics, we identified nine surface-exposed amino acid residues that are critical for the DNA-binding activity of MipZ. To verify the direct involvement of these residues in the interaction with DNA, we mapped the DNA-binding interface of MipZ by hydrogen/deuterium exchange mass spectrometry (HDX-MS), a method detecting local shifts in the accessibility of backbone amide hydrogens caused by ligand binding (42). To this end, the dimeric D42A variant was transferred into deuterated buffer and incubated with or without a short (14 bp) dsDNA oligonucleotide. After fragmentation and mass spectrometric analysis of the protein, we then searched for peptides that showed a reduced rate of deuterium uptake in the presence of DNA. A total of 168 peptides were unambiguously identified, covering 94% of the protein sequence. Protected peptides mapped to three distinct regions (R1, R3, and R4) of MipZ (**Fig. 4A-B and S7A**). A closer inspection of the data showed that R1 corresponds to the Walker A motif, suggesting a conformational change in this region upon DNA binding. R3 is only partially exposed on the dimer surface, and mutations in this region (D147A, T150A, E152A) do not affect the biological function of MipZ (**Fig. S8**). This region may therefore be subject to conformational changes upon DNA binding but not directly contribute to the interaction. The elements constituting R4, by contrast, are located on the dimer surface and contain all of the residues identified in the mutant screen (**Fig. 4B**), confirming the relevance of the *in vivo* results. Together, these findings support the hypothesis that the DNA-binding interface of MipZ is formed by a linear array of positively charged amino acids (**Fig. 4C**) that mediate non-specific DNA binding through interaction with the DNA phosphate backbone.

**Figure 4.**
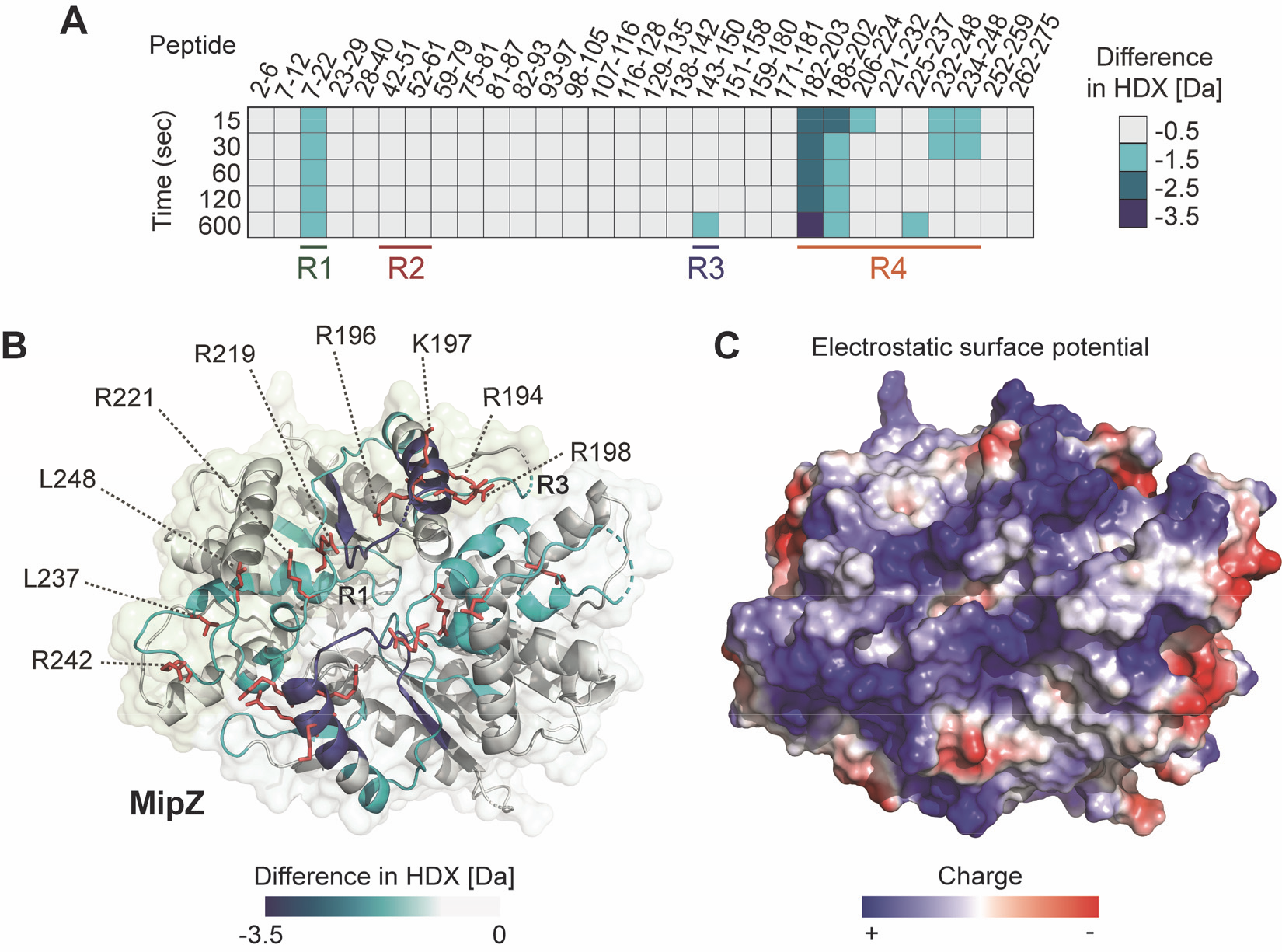
Hydrogen/deuterium exchange (HDX) analysis of the MipZ-DNA interaction. (**A**) Heat plot listing the maximal differences in deuterium uptake between the DNA∙MipZ-D42A complex and MipZ-D42A alone at different incubation times for a series of representatives peptides. The color code is given on the right. All experiments were performed in the presence of ATPγS to further increase the stability of the dimer. The four different regions that are protected upon DNA binding to MipZ or ParA (see **Figure 5**) are indicated at the bottom. (**B**) Mapping of the maximum differences in HDX observed upon DNA binding onto the crystal structure of the MipZ dimer (PDB ID: 2XJ9). The color code is given at the bottom. The DNA-binding residues identified in the mutant screen are shown in red. A surface representation of the structure is shown in the background. For clarity, the ATP and Mg^2+^ ligands are not shown. (**C**) Electrostatic surface potential of the MipZ dimer. The color code is given at the bottom.

### MipZ and ParA share a similar DNA-binding region

MipZ and ParA share the ability to interact non-specifically with DNA. Previously, two arginine residues at the dimer interface of ParA were shown to be critical for DNA binding (28, 35). Moreover, a recent crystallographic study has suggested that *Hp*ParA interacts with DNA in a region homologous to the DNA-binding interface of MipZ (37). However, a comprehensive analysis of the determinants that mediate the ParA-DNA interaction in solution is still missing. To address this issue, we extended our HDX studies to the ParA homolog of *C. crescentus* and mapped the regions of the protein that were protected upon interaction with a dsDNA oligonucleotide (**Fig. 5A**). Since it was not possible to purify an ATPase-deficient (D44A) variant of ParA in soluble form, the experiments were performed with the wild-type protein, locked in the dimeric state by addition of ATPγS. In total, 78 peptides were unambiguously identified, which covered 94% of the complete protein sequence. Interestingly, out of the three protected regions, two (R1 and R4) were homologous to regions protected upon DNA binding in MipZ, whereas one (R2) was specifically detected in ParA (**Fig. 5A-B and S7B**). The elements constituting R2 are mostly buried in the interior of the ParA dimer, suggesting that they do not directly contribute to DNA binding. Notably, however, they also include a loop with an arginine residue (R61) that is exposed on the lateral side of the complex. As in MipZ, R4 showed the highest degree of protection. It stretches along the edge of the dimer interface and includes six positively charged and two bulky hydrophobic residues that are located at the protein surface (**Fig. 5B-C**). Mutational analysis confirmed that all of these residues are critical for proper ParA localization and function *in vivo* (**Fig. S9**). In line with previous work (35, 37), these findings demonstrate that the DNA-binding activity of ParA also relies on non-specific electrostatic interactions with the DNA phosphate backbone. Interestingly, however, only one of the residues identified in ParA (K233) has an equivalent in the DNA-binding pocket of MipZ (R242), because the DNA-interacting region of MipZ is largely part an insertion that is not conserved in other P-loop ATPases (**Fig. S10**). The DNA-binding pockets of the two proteins may thus have evolved independently, even though the principal mechanism of target recognition is identical in both cases.

**Figure 5.**
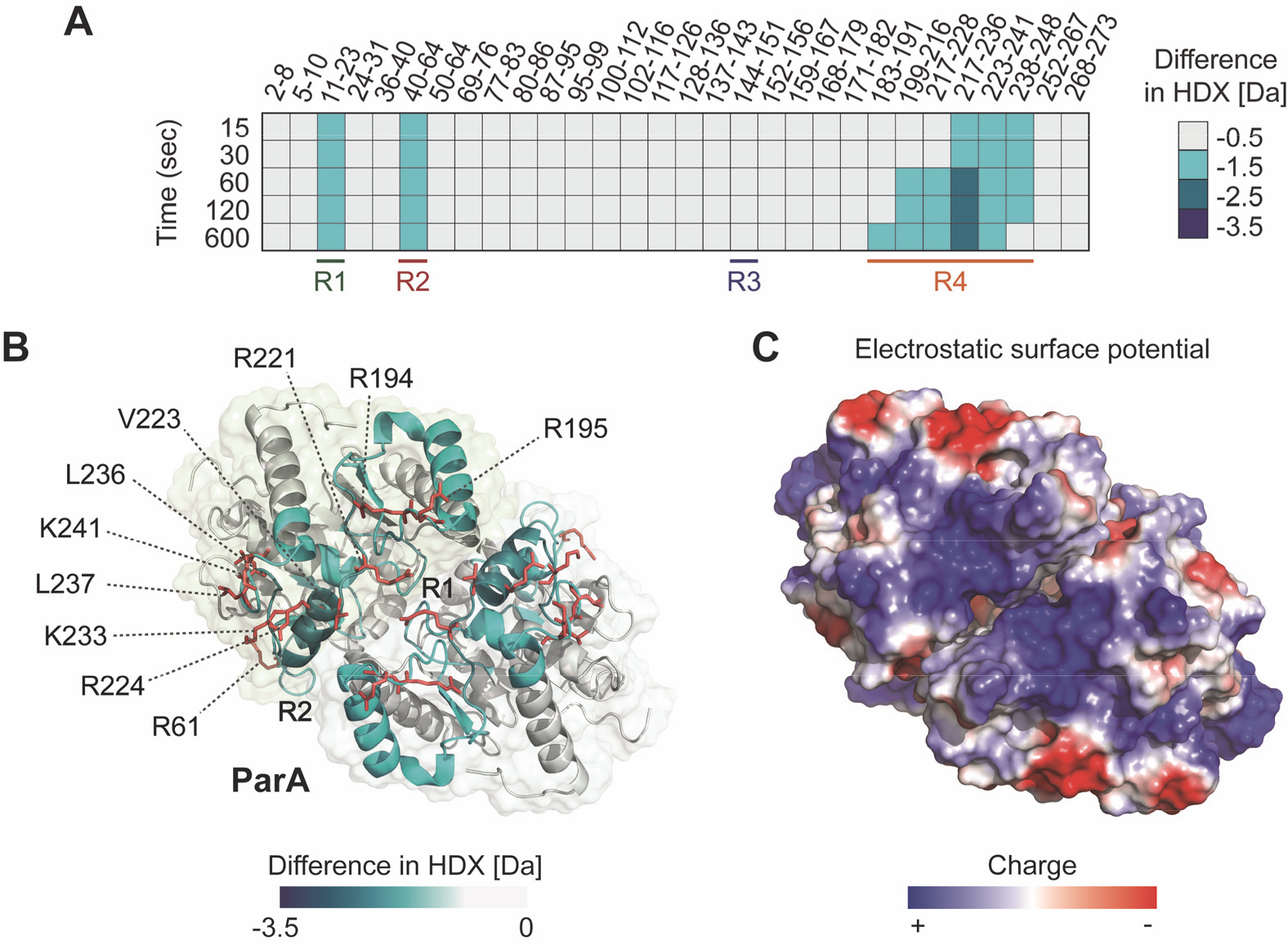
Hydrogen/deuterium exchange (HDX) analysis of the *C. crescentus* ParA-DNA interaction. (**A**) Heat plot listing the maximal differences in deuterium uptake between the DNA∙ParA complex and MipZ alone at different incubation times for a series of representatives peptides. The color code is given on the right. All experiments were performed in the presence of ATPγS to lock the protein in the dimeric state. The four different regions that are protected upon DNA binding to MipZ or ParA (see **Figure 4**) are indicated at the bottom. (**B**) Mapping of the maximum differences in HDX observed upon DNA binding onto a structural model of ParA, generated with *Hp*ParA (PDB ID: 6IUB) (37) as a template. A surface representation of the structure is shown in the background. For clarity, the ATP and Mg^2+^ ligands are not shown. (**C**) Electrostatic surface potential of the ParA dimer. The color code is given at the bottom.

## DISCUSSION

In this work, we used a combination of biochemical and genetic approaches to identify and comprehensively analyze the DNA-binding sites of the P-loop ATPases MipZ and ParA from *C. crescentus*. Our results show that both proteins associate with DNA in a sequence-independent manner through electrostatic interactions with the phosphate backbone (**Fig. 6**). However, despite this similarity in the mechanism of action, the architecture of the binding sites varies considerably. Interestingly, most P-loop ATPases studied to date share the DNA-binding region of ParA (4, 20, 35–37) (**Fig. S10**). The interaction determinants of MipZ, by contrast, are largely located in a segment of the protein that is not conserved in other family members. This finding supports the idea that the ability of MipZ to interact with DNA may have evolved independently. Alternatively, its DNA-binding interface may have changed to modulate the kinetics of the interaction or the effect of DNA on the ATPase cycle or function of MipZ.

**Figure 6.**
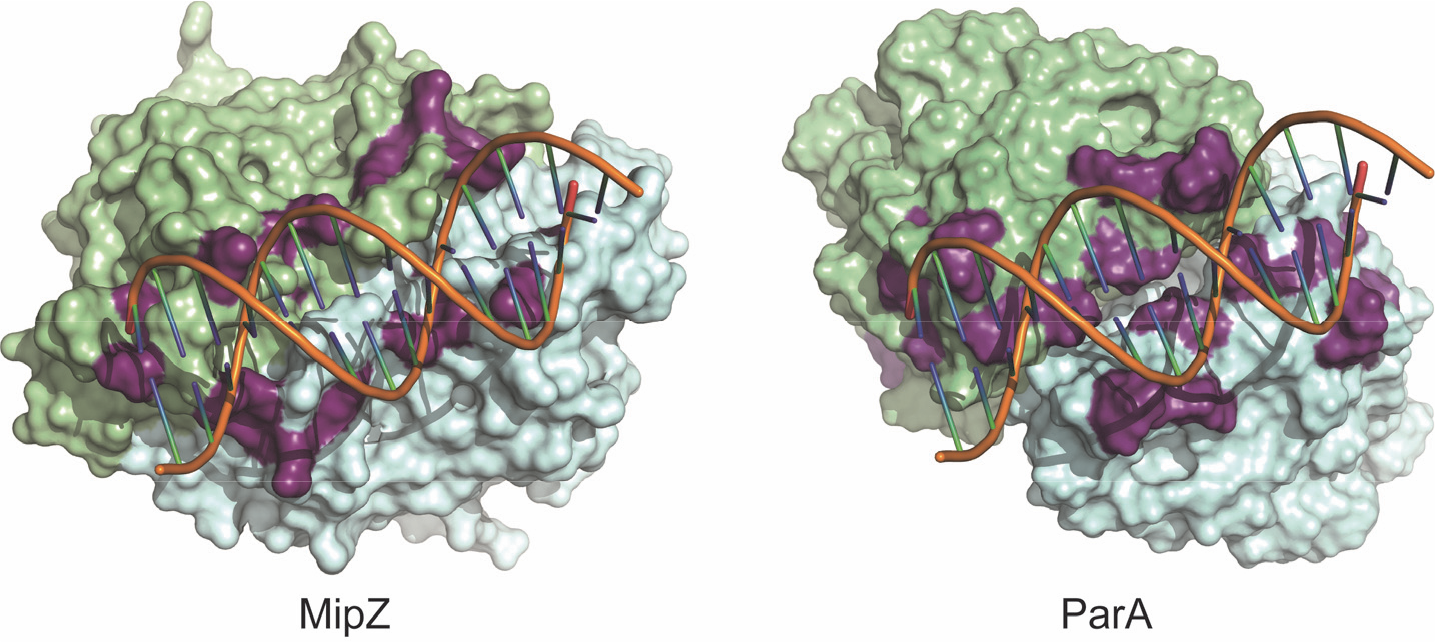
Comparison of the DNA-binding regions of MipZ and ParA from *C. crescentus*. Surface representations of the dimer structures of (**A**) MipZ (PDB ID: 2XJ9) and (**B**) ParA (modeled as described in **Fig. 5B**). Residues shown to be important for DNA binding are highlighted in purple. Short dsDNA oligonucleotides were modeled into the experimentally verified DNA-binding interfaces. The two subunits of each dimer are depicted in green and blue, respectively.

Our study identified a total of nine residues that critically contribute to DNA binding by MipZ (**Fig. 4**). The individual importance of these residues appears to differ significantly. Despite their peripheral location in the DNA-binding site, R194 and R198 are essential for interaction with DNA, consistent with the observation that they are part of a region that is highly protected from the environment upon DNA binding. The remaining residues make additional contacts that contribute to stabilizing the MipZ-DNA interaction. Interestingly, while the majority of them are positively charged and, thus, suited to directly contact the DNA phosphate backbone, two (L237 and L248) had bulky hydrophobic side chains. Mutations in these leucines had only moderate effects on the functionality of MipZ *in vivo*, although the R237A variant showed a strongly reduced DNA-binding activity *in vitro*. The precise role of these residues remains to be investigated. Notably, the crystal structure of the *H. pylori* ParA-DNA complex revealed a conserved leucine residue that is in direct contact with the DNA molecule. However, this interaction involves the main chain amide group rather than the hydrophobic side chain (37). A similar situation may be observed for MipZ, with mutations in L237 and L248 leading to distortions in the main chain that impede its interaction with the phosphate backbone. Interestingly, DNA binding also leads to conformational changes in the Walker A motif, thus likely affecting the kinetics of DNA hydrolysis. Consistent with this idea, previous work has shown a stimulatory effect of DNA on the ATPase activity of MipZ (13). Since the DNA-binding site of MipZ spans the dimer interface and comprises residues from both subunits, only the dimeric complex should have full DNA-binding activity. This hypothesis is supported by the observation that monomeric MipZ variants are unable to interact with DNA *in vitro* or stably associate with the nucleoid *in vivo* (13) (**Fig. 2**). Notably, a comparison of the MipZ monomer and dimer structures shows that the DNA-binding regions undergo conformational changes upon dimerization, which may contribute to stabilizing the MipZ-DNA interaction (**Fig. S11**). In line with the lack of target specificity *in vitro*, ChIP-seq experiments demonstrated that MipZ does not bind to distinct chromosomal sites *in vivo*. Nevertheless, it is not evenly distributed across the nucleoid but enriched in GC-rich regions, potentially due to the influence of competing proteins that preferentially interact with AT-rich sequences. It will be interesting to see whether other ParA-like ATPases with non-specific DNA-binding activity (20, 35, 37, 38) show a similar bias in their distribution.

Our analysis of the DNA-binding determinants of ParA identified ten positively charged or bulky hydrophobic residues (**Fig. 5**) that are located in regions protected by DNA *in vitro* and critical for ParA function *in vivo*. Two of these residues (R195 and R224) are conserved in several other ParA-like ATPases (**Fig. S10**), and one or both of their counterparts have previously been implicated in the DNA-binding activity of other ParA homologs (28, 35, 37), PpfA (20), and PomZ (17). Moreover, the residues equivalent to R221, V223, R224 and K241 are in direct contact with DNA in the crystal structure of *Hp*ParA (37), indicating an excellent agreement between the HDX and crystallographic data. In addition, we identified three more residues that are critical for DNA binding by *Cc*ParA (K233, L236, L237) and also conserved in other ParA homologs, suggesting that they could be core components of the DNA-binding site in members of the ParA family. By contrast, R194 of *Cc*ParA is not conserved in other ParA homologs but present in ParC. In general, a comparison of different ParA-like ATPases shows that the individual constituents of their DNA-binding sites are often poorly conserved (**Fig. S10**). However, most crystallized members of this family share a patch of positively charged residues that stretches along the dimer interface in corresponding regions of the protein (**Fig. S12**). Modeling studies with *Hp*ParA as a template suggest that the same is true for other ParA-like ATPases, such as PpfA, ParC, VirC1 and PomZ, whose structures still remain to be solved. A notable exception to this rule is the cell division regulator MinD, which was shown to interact non-specifically with DNA and contribute to chromosome segregation in *E. coli* (43). In this protein, the region that typically forms the DNA-binding interface shows a high density of negative charges and only contains a single arginine residue (R219), which is essential for DNA binding (43). These observations suggest that MinD may use a different mode of DNA binding than its family relatives. Interestingly, a recent crystal structure of the archaeal plasmid pNOB8 ParA-DNA complex revealed a unique mode of DNA-binding in which the ParA dimer interacts laterally with two DNA molecules. In doing so, it undergoes a large conformational change, with one subunits rotating ~38° relative to the second subunit, thereby opening the active site and exposing the sandwiched nucleotides to the solvent (38). Our HDX analysis of *Cc*ParA and MipZ did not reveal any obvious shift in the accessibility of the dimer interface upon DNA binding. Moreover, both proteins displayed only a single DNA-binding site per dimer. In line with previous studies (35, 37), we therefore suggest that ParA-like ATPases typically have similar conformations in the DNA-free and complexed state and bind DNA through a single patch of positively charged residues located at the edge of the dimer interface. Collectively, our results support the notion that ParA-like ATPases represent a diverse family of regulators that share common functional principles but have diverged considerably to adapt to distinct cellular functions.

## EXPERIMENTAL PROCEDURES

### Bacterial strains, plasmids and growth conditions

The bacterial strains, plasmids and oligonucleotides used in this work are listed in **Tables S1-S3**. *C. crescentus* CB15N and its derivatives were cultivated in PYE (peptone-yeast-extract) medium at 28 °C, supplemented with antibiotics when appropriate at the following concentrations (μg ml^−1^; liquid/solid medium): kanamycin (5/25), streptomycin (5/5), spectinomycin (25/50). To induce the expression of genes placed under the control of the Pxyl or Pvan promoters, media were supplemented with 0.3% (w/v) D-xylose or 0.5 mM sodium vanillate, respectively. *E. coli* TOP10 (Invitrogen, USA) was used for cloning purposes. Proteins were overproduced in Rosetta (DE3)pLysS (Novagen, Germany). *E. coli* cells were cultivated aerobically in Luria-Bertani broth at 37 °C, supplemented when appropriate with 0.5% (w/v) glucose and antibiotics at the following concentrations (μg ml^−1^; liquid/solid medium): ampicillin (200/200) or chloramphenicol (20/30). Protein overproduction was induced by addition of 1 mM isopropyl-β-D-thiogalactopyranoside (IPTG).

### Light and fluorescence microscopy

Cells were immobilized on 1 % agarose pads and imaged using an Axio Imager.M1 microscope (Carl Zeiss AG, Germany) equipped with a Photometrics Cascade:1K EMCCD camera or a Zeiss Axio.Imager Z1 microscope equipped with a pco.edge 4.2 sCMOS camera (PCO). Images were acquired with a Zeiss Plan-Apochromat 100×/1.40 Oil Ph3 M27 objective. An X-Cite^®^120PC metal halide light source (EXFO, Canada) and an ET-YFP filter cube (Chroma, USA) were used for fluorescence imaging. Imaging data were analyzed with Metamorph 7.7 (Molecular Devices, USA) or Fiji 1.49 (44). Violin and boxplots were generated with R version 3.5.1 (http://www.r-project.org). The quantification of gradient patterns was performed in MATLAB R2014b (Mathworks, USA). To generate demographs, fluorescence intensity profiles were measured with Fiji and processed in R using the Cell Profiles script (45). Details of the FRAP analysis are provided in Supplemental Experimental Procedures.

### Protein purification and*in vitro* assays

The purification of proteins and the different *in vitro* assays used in this study are described in the Supplemental Experimental Procedures. Immunodetection was performed according to standard protocols using a polyclonal anti-MipZ antiserum (1:10,000) (6) or an affinity-purified polyclonal anti-GFP antibody (1:10,000) (#G1544; Sigma-Aldrich, Germany). Immunocomplexes were detected with a secondary goat anti-rabbit antibody conjugated with horseradish peroxidase and visualized with the Western Lightning plus–ECL chemiluminescent reagent (PerkinElmer, USA) in a ChemiDoc MP imaging system (Bio-Rad Laboratories, USA). Images were aquired in ImageLab 5.0 (Bio-Rad Laboratories, USA) and processed in Adobe Illustrator CS5 (Adobe Systems, USA).

## ACKNOWLEDGEMENTS

We thank Daniela Kiekebusch, Till Ringel and Jaspara Knopp for help with the alanine-scanning mutagenesis, and Luis M. Oviedo for assistance with MatLab. This work was funded by the German Research Foundation (DFG) (project 269423233 – TRR 174; to G.B. and M.T.), the Swiss National Science Foundation (grant 31003A_182576; to P.H.V) and the Max Planck Society (Max Planck Fellowship to M.T.). We acknowledge support from the DFG-Core Facility for Interactions, Dynamics and Macromolecular Assembly Structure (MIDAS) (to G.B.). L.C.-G. received funding from the Horizon Research and Innovation Program of the European Union (Grant Agreement 659174; MIPZ).

## AUTHOR CONTRIBUTIONS

B.H. generated strains and conduced *in vivo* and *in vitro* studies. Y.R. purified the proteins and performed the biolayer interferometry analysis. L.C.-G. generated strains, performed *in vivo* analyses and conducted the FRAP experiments. G.P. performed the ChIP-Seq analysis. W.S. performed the HDX experiments. B.H., Y.R., L.C.-G., W.S. and M.T. analyzed the data. P.V., G.B. and M.T. supervised the work. Y.R., L.C.-G. and M.T. wrote the paper, with input from all other authors.

